# Glycoprofiling of proteins as prostate cancer biomarkers: a multinational population study

**DOI:** 10.1101/2023.06.27.546717

**Authors:** Andrea Pinkeova, Adela Tomikova, Aniko Bertokova, Eva Fabinyova, Radka Bartova, Eduard Jane, Stefania Hroncekova, Karl-Dietrich Sievert, Roman Sokol, Michal Jirasko, Radek Kucera, Iris E. Eder, Wolfgang Horninger, Helmut Klocker, Petra Ďubjaková, Juraj Fillo, Tomas Bertok, Jan Tkac

## Abstract

The glycoprofiling of two proteins, the free form of the prostate-specific antigen (fPSA) and zinc-α-2-glycoprotein (ZA2G), was assessed to determine their suitability as prostate cancer (PCa) biomarkers. The glycoprofiling of proteins was performed by analysing changes in the glycan composition on fPSA and ZA2G using lectins (proteins recognising glycans, *i.e*. complex carbohydrates). The specific glycoprofiling of the proteins was performed using magnetic beads (MBs) modified with horseradish peroxidase (HRP) and antibodies that selectively enriched fPSA or ZA2G from human serum samples. Subsequently, the antibody-captured glycoproteins were incubated on lectin-coated ELISA plates. In addition, a novel glycoprotein standard (GPS) was used to calibrate the assay. The glycoprofiling of fPSA and ZA2G was performed in human serum samples obtained from men undergoing prostate biopsy after an elevated serum PSA, and prostate cancer patients with or without prior therapy. The results are presented in the form of a ROC (Receiver Operating Curve). A DCA (Decision Curve Analysis) to evaluate the clinical performance and net benefit of fPSA glycan-based biomarkers was also performed. While the glycoprofiling of ZA2G showed little promise as a potential PCa biomarker, the glycoprofiling of fPSA would appear to have significant clinical potential. Hence, the GIA (Glycobiopsy ImmunoAssay) test integrates the glycoprofiling of fPSA (*i.e*. two glycan forms of fPSA). The GIA test could be used for early diagnoses of PCa (AUC=0.84; n=501 samples) with a potential for use in therapy-monitoring (AUC=0.85; n=168 samples). Moreover, the analysis of a subset of serum samples (n=215) revealed that the GIA test (AUC=0.81) outperformed the PHI (Prostate Health Index) test (AUC=0.69) in discriminating between men with prostate cancer and those with benign serum PSA elevation.

## 1. Introduction

In 2020, the worldwide incidence of and mortality from prostate cancer (PCa) were estimated as 1.41 million and 375,000, respectively; these figures are predicted to increase to 2.24 million and 721,000 by 2040 [1]. PCa has a large impact on a patient’s quality of life; it significantly influences sexual, bowel and urinary functions [2]. Early detection of cancer is crucial for a chance of curative treatment; however, PCa screening also identifies PCa cases that are not fatal, thereby causing significant social distress, or leading to unnecessary subsequent overtreatment [2]. Detection of indolent, low-risk PCa (*i.e*. Gleason score 3 + 3 or ISUP grade group 1) may lead to anxiety and depression, especially for patients subsequently undergoing active surveillance [3]. Any method yielding reliable information about the presence and grade of tumours in biopsy-naïve patients (so-called liquid biopsy methods) may prevent overdiagnosis and overtreatment, and increase the quality of life of patients, especially those suffering from clinically insignificant low-risk PCa [4].

Re-designing screening and diagnostic programmes that benefit patients and implementing novel, non-invasive procedures with reduced or no side-effects are very important. Novel PCa biomarkers are actively sought so as to improve patient management, reduce the number of negative biopsies and, thereby, healthcare system expenses and, importantly, lower future barriers between clinicians and asymptomatic patients [5]. In recent years, many different types of biomolecules have been proposed as PCa biomarkers: small molecules/metabolites, nucleic acids (including miRNAs, mRNA, circulating tumour DNA), proteins, extracellular vesicles (exosomes) and circulating tumour cells [6-10]. Post-translational modifications of proteins, especially glycosylation, were shown to be strongly associated with disease development and progression [11, 12].

The glycosylation process takes place in the Golgi apparatus, an organelle continuously receiving and processing a flow of protein cargoes. Its well-organised cisternal structure has been shown to be crucial for its proper functioning. Oncogenesis disrupts the structural integrity of the Golgi apparatus, resulting in the abnormal expression of enzymes, the dysregulation of anti-apoptotic kinases and the hyperactivity of myosin motor proteins [13]. Moreover, the structural alterations and fragmentation of the Golgi apparatus during oncogenesis lead to the aberrant glycosylation of proteins: for example, sialylation associated with epithelial-mesenchymal transition and extracellular matrix remodelling [14, 15]. These altered glycosylation patterns result from changed activity of glycosyltransferases. Since glycosyltransferases are anchored into the Golgi apparatus membrane, their activity is influenced by the structural remodelling of the Golgi apparatus [16, 17].

In previous studies, we demonstrated the diagnostic potential of glycosylation changes in free prostate-specific antigen (fPSA). Aberrant sialylation and fucosylation, for example, can be used to diagnose both early-stage PCa and high-grade prostatic intraepithelial neoplasia; they can even be used in the recognition of a castration-resistant form of PCa [18, 19]. In our Glycobiopsy Immuno Assay (GIA) test, we use a unique magnetic-beads-based protocol that overcomes the challenges inherent in lectin-assisted glycoprofiling of proteins [20-22]. The magnetic beads are modified by anti-fPSA antibodies for the selective enrichment of fPSA from human blood serum samples. Subsequently, the magnetic beads with attached fPSA are added to lectin-coated ELISA plates in order to perform glycoprofiling. Finally, the sandwich ELISA protocol is completed by a horseradish peroxidase (HRP) reaction, (see detailed protocol on: www.glycanostics.com). This protocol has proved to be robust and reproducible.

In the aforementioned studies [18, 19], one serum sample of one particular PCa patient was applied to calibrate the analysis and correct for plate-to-plate variability. Such an approach was feasible for clinical validation using only a limited number of samples and/or for the analysis of samples in a single run/day. The analysis of a large set of samples, or the analysis over a longer period of time, requires a proper calibration. Attempts to resolve this issue by producing fPSA with attached cancer-specific glycans in cultured cancerous prostate cell lines were not successful because the glycans present on cell-line derived-fPSA differ significantly from those glycans present on fPSA collected from PCa patients [23]. In addition, commercially available fPSA is not suitable for calibration since it is isolated from healthy individuals/donors [24, 25] and does not contain cancer-specific glycans. This issue was resolved by developing a glycoprotein standard (GPS) - streptavidin protein with chemically attached glycans - to calibrate the GIA test [26]. GPS calibration was an integral part of the current clinical study and, to our knowledge, this is the first glycoprofiling study using this new approach.

In the present study, serum samples from Caucasian men from four different European countries (the Slovak Republic, the Czech Republic, Austria and Germany) were analysed with the objective of determining whether the GIA test could be applied to diagnostics and therapy monitoring. The study sought to compare the clinical performance of the GIA test to the performance of serological tests based on an analysis of PSA forms such as tPSA, fPSA and a combination thereof (Prostate Health Index (PHI) detecting tPSA (total PSA), fPSA (free form of PSA) and −2proPSA isoforms). In addition, we investigated the glycoprofiling of zinc-α-2-glycoprotein (ZA2G) to increase the overall accuracy of glycan-based PCa diagnostics. A ROC (Receiver Operating Curve) analysis and a DCA (Decision Curve Analysis) were used as two independent statistical methods to evaluate the benefit of these glycan-based assays in clinical practice.

## 2. Materials and Methods

### 2.1. Clinical samples

The serum samples used in the study were taken from (i) the Department of Urology, Medical University Innsbruck, Austria (serum samples present in the biobank collected up to 10/2016 were used), (ii) Klinikum Lippe - Clinic for Urology in Detmold, Germany (serum samples collected in the period 10/2020 – 01/2021), (iii) University Hospital in Pilsen, the Czech Republic (serum samples collected in the period 06/2021 – 02/2022) and (iv) Private Urological Ambulance in Trencin, the Slovak Republic (serum samples collected in the period 05/2021 – 02/2022). All the men underwent a prostate transrectal ultrasound-guided prostate biopsy after presenting with elevated serum tPSA. The clinical characteristics of the participants whose samples were used in the study are summarised in **Table 1**. The authors did not have access to information that could identify individual participants during or after data collection.

**Table 1:**
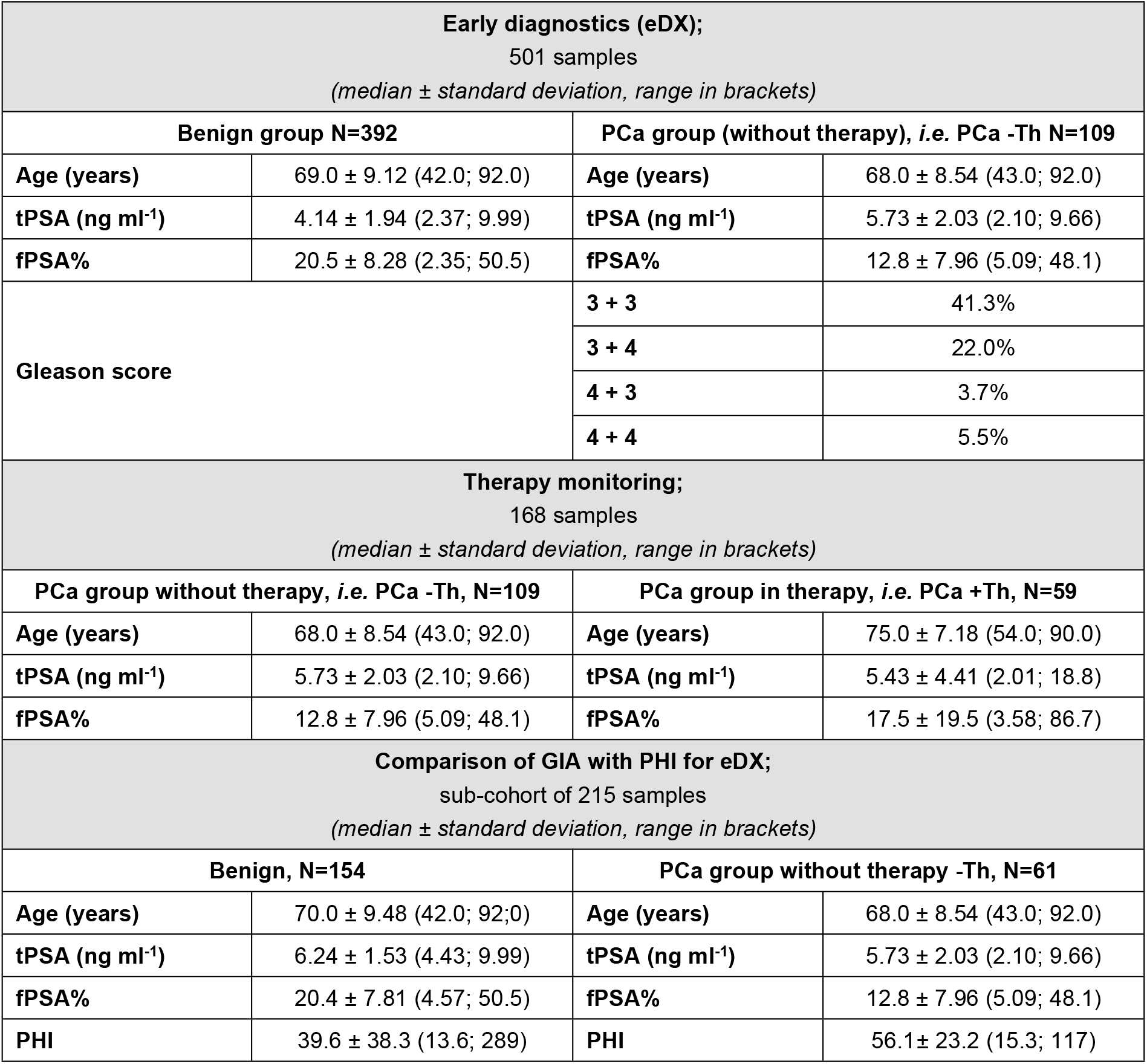
Clinical characteristics of cohorts applied to early diagnostics (eDX), therapy monitoring, and GIA to PHI test comparison analyses.

All the samples were collected *prior* to radical prostatectomy and the study was reviewed and approved by the respective Ethics Committees (Eticka komisia Trenčianskeho samosprávneho kraja, Trenčín, Slovakia; Etická komise FN a LF UK v Plzni, Plzeň, Czech Republic; Ethikkommision der Medizinischen Universität Innsbruck, Innsbruck, Austria; and Ethik-Kommission Westfalen-Lippe, Munster, Germany) with written consent obtained. Based on the biopsy results, a cancer cohort and a benign cohort were chosen; both cohorts fulfil the criteria of a “grey zone” with serological tPSA levels in the ranges of 2-10 ng mL^-1^; the cohorts were also similar in age and tPSA levels. The PCa cohort was subdivided into low-risk (Gleason score 3+3, ISUP GG 1) and high-risk PCa (Gleason score ≥ 7, ISUP GG ≥2) sub-groups based on histological examinations of biopsied tissues.

Two independent clinical validation studies were performed; namely, (i) early diagnostics (early DX; benign *vs*. PCa, no *prior* therapy) using 501 samples (unless indicated otherwise, see **Table 1**) and (ii) therapy monitoring (PCa with no *prior* therapy *vs*. PCa with *prior* therapy of any kind) using 168 samples from PCa patients. A comparison of the GIA test with the Prostate Health Index (PHI from Beckman Coulter) was performed on a subgroup of 215 benign and PCa samples for which PHI values were measured.

### 2.2. Analyses and biostatistics

The glycoprofiling of proteins was performed using WFL (the *Wisteria floribunda* agglutinin that recognises *N*-acetylgalactosamine, *i.e*. GalNAc and *N*-acetygalactosamine linked to *N*-acetylglucosamine structures, *i.e*. LacdiNAc) and PHA-E (the *Phaseolus vulgaris* erythroagglutinin that recognises more complex structures, *i.e. N*-glycans with outer galactose, *i.e*. Gal and bisecting *N*-acetylglucosamine, *i.e*. GlcNAc) [27], as published previously [18, 19]. The results from the glycoprofiling of fPSA by two lectins (WFL and PHA-E) were obtained as fPSA^WFL^ and fPSA^PHA-E^ values. The GIA test was evaluated by application of the newly developed Glycoprotein Standard (GPS, glycosylated streptavidin). A detailed description of the test and the method for preparation of the standard is provided in the supporting information. Both lectins in their unconjugated form were purchased from Vector Labs (USA). The anti-fPSA antibody and all the anti-ZA2G antibodies were purchased from Abcam (UK). Streptavidin was purchased from Vector Labs, USA and the anti-streptavidin antibody from MyBioSource (USA). Other common chemicals and buffer components were purchased from Sigma-Merck (USA).

## 3. Results

### 3.1. Early diagnostics of PCa using protein glycoprofiling

#### 3.1.1. GIA test

Reducing the number of negative biopsies by increasing the accuracy of screening and diagnostic methods remains an unmet medical need. The GIA test overcomes the common disadvantages of lectin biorecognition; *i.e*. weak ligand-receptor interactions and a lack of substrate specificity. The GIA test was calibrated using GPS, the preparation of which and use for calibration are detailed in the Supporting Information file. The method was used for discriminating between prostate cancer and prostate benign serum samples. The 392 benign and 109 PCa serum sample values were subjected to ROC curve analysis and yielded an excellent AUC of 0.84 (**Fig. 1**). At 95% specificity the sensitivity was 40.4%, while at 95% sensitivity the specificity was 38.0%. The confidence interval CI (95%) for the AUC values was [0.79 - 0.88]. In comparison, tPSA provided an AUC of 0.68 (CI (95%) [0.62 – 0.73]). At 95% specificity the sensitivity was 11.9%, while at 95% sensitivity the specificity was 4.8%. The fPSA analysis provided an AUC of 0.60 (CI (95%) [0.62 – 0.73]). At 95% specificity the sensitivity was 9.2%, while at 95% sensitivity the specificity was 11.5%. The fPSA% analysis revealed an AUC of 0.76 (CI (95%) [0.71 – 0.81]). At 95% specificity the sensitivity was 25.3%, while at 95% sensitivity the specificity was 22.9%. A combination of biomarkers tPSA and fPSA provided an AUC of 0.78 (CI (95%) [0.73 – 0.83]), a value only slightly higher than the AUC value for fPSA% of 0.76. At 95% specificity the sensitivity was 63.3%, while at 95% sensitivity the specificity was 25.3%. A combination of biomarkers tPSA and fPSA% provided an AUC of 0.78 (CI (95%) [0.73 – 0.83]). At 95% specificity the sensitivity was 36.7% while at 95% sensitivity the specificity was 74.8%. In this comparison, GIA outperformed all PSA and PSA combination parameters in discriminating between patients with cancer and benign prostate histology biopsy results.

**Figure 1:**
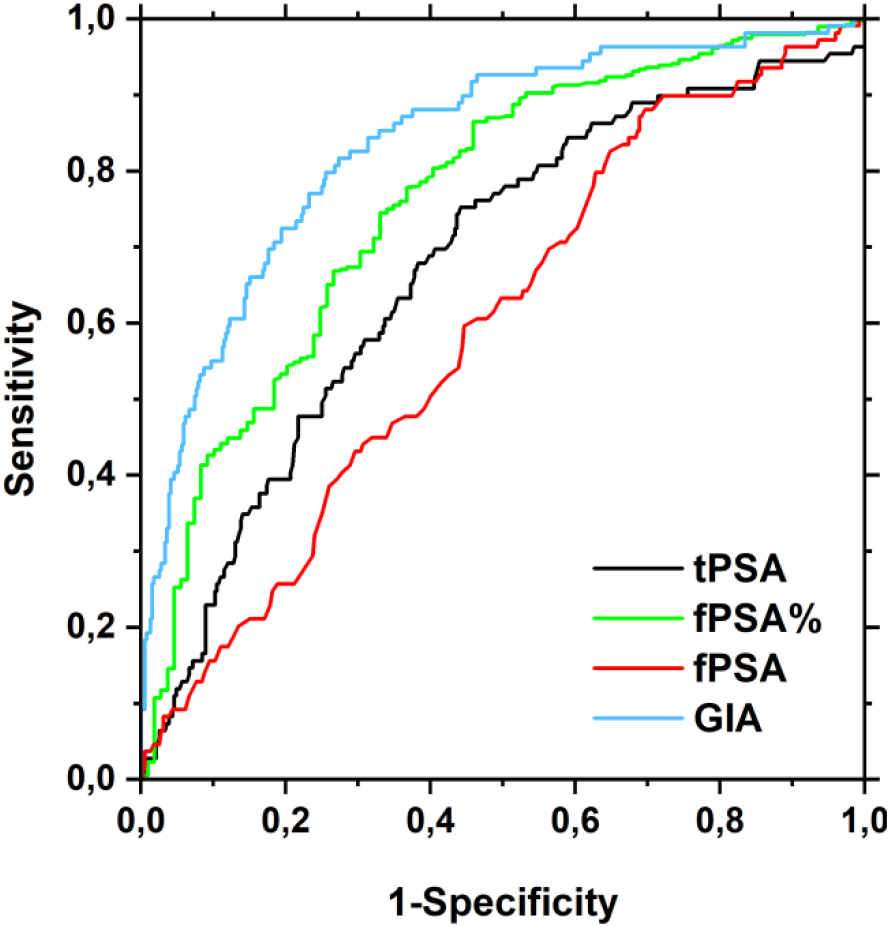
ROC analysis depicting tPSA (black line), fPSA (red line), fPSA% (green line) and GIA test (blue line) curves for cases of early DX (501 samples in total). The AUC values are 0.68 and 0.84 for tPSA and GIA test, respectively.

In a subgroup analysis, the application of the GIA test to the identification of low-risk and high-risk PCa patients was investigated. Serum samples from 47 PCa individuals with low-risk (Gleason score 3 + 3, ISUP GG 1) and 41 high-risk PCa (Gleason score ≥ 7, ISUP GG ≥2) were compared. The GIA test yielded a higher AUC value (0.67) than the tPSA test (0.57) and than the combination of tPSA+fPSA (0.64).

A calculation of the number of negative (avoidable) biopsies identified by the PCa biomarkers (tPSA, fPSA, fPSA% and GIA test at 80% sensitivity revealed the following percentages: tPSA 30%, fPSA 53%, fPSA% 53% and GIA test 70% (**Fig. 2**). Hence, the GIA test has the potential to significantly reduce the number of negative (avoidable) biopsies, whereas the tPSA and fPSA tests have a limited potential. While the GIA test missed only six high-risk PCa cases (Gleason score ≥ 7, ISUP GG ≥2), tPSA missed eight high-risk PCa cases, fPSA missed seven high-risk PCa cases and fPSA% missed five high-risk PCa cases.

**Figure 2:**
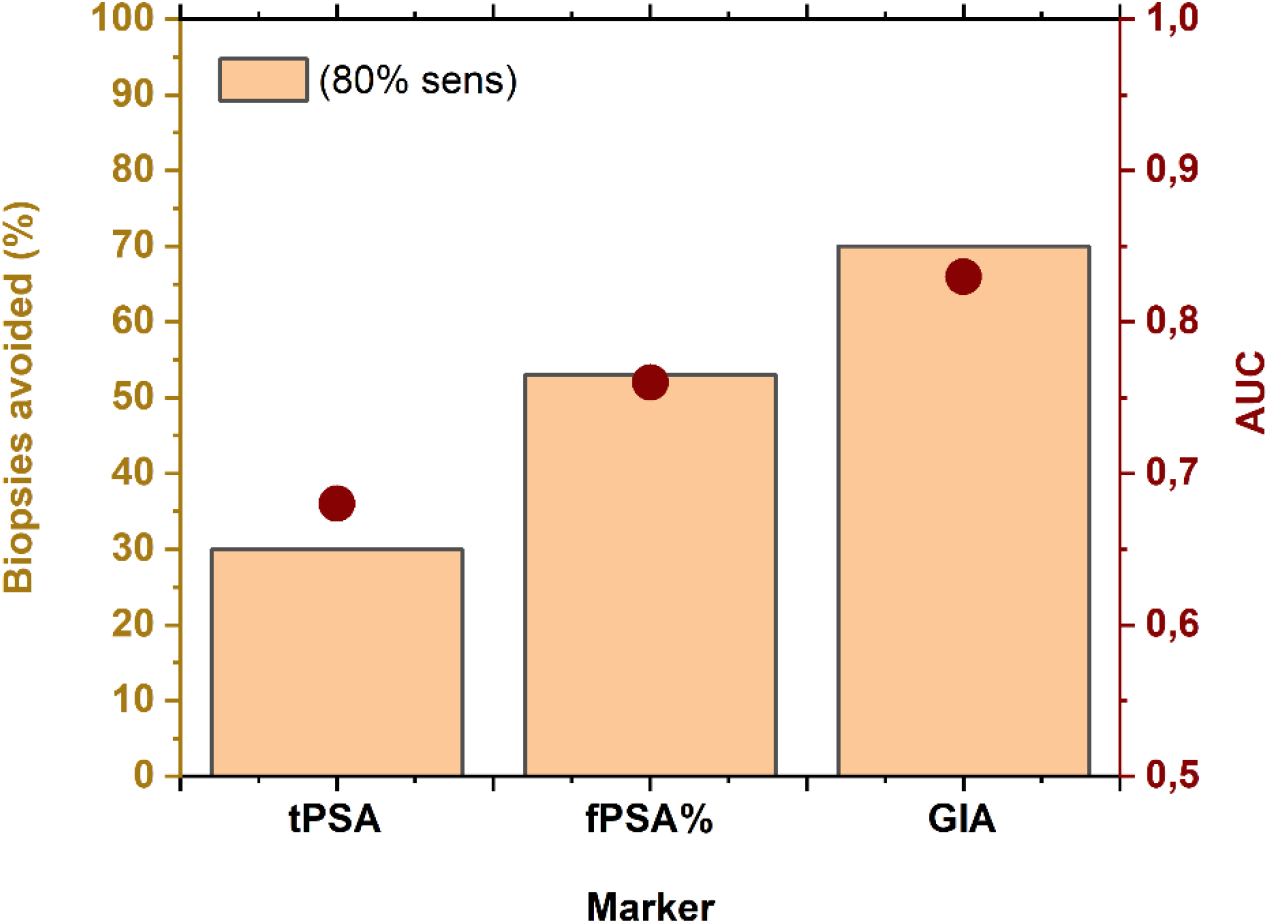
Percentage of negative (avoidable) biopsies (orange columns) calculated at 80% sensitivity for PCa biomarkers tPSA, fPSA and GIA test, respectively; results are based on 501 serum samples. Negative (avoidable) biopsies were calculated as the ratio of correctly identified benign patients from among the whole benign cohort.

The effect of age (<50, 50-60, 60-70 and >70) on correct PCa diagnostics by the GIA test was evaluated. Out of the 501 samples involved in the early diagnosis of PCa evaluation, twelve samples from individuals who were younger than 50 years yielded an AUC of 1.00 (due to the small cohort), 108 individuals from individuals aged 50-60 years an AUC of 0.87 ± 0.15, 160 individuals aged 60-70 years an AUC of 0.87 ± 0.12 and the largest age group of 221 individuals older than 70 years an AUC value of 0.81 ± 0.14, suggesting a consistent AUC value obtained across different age groups.

#### 3.1.2. Glycoprofiling of ZA2G

The rationale behind the glycoprofiling of ZA2G was to increase the accuracy of PCa diagnostics by a combination of more biomarkers, *i.e*. glycoprofiling of fPSA with glycoprofiling of ZAG2. Integration of the glycoprofiling of ZA2G using PHA-E and WFL lectins (*i.e*. ZAG2^PHA-E^ and ZAG2^WFL^) yielded a very low AUC of (0.52 and 0.54, respectively) in the discrimination between malignant and benign patients. In comparison, the glycoprofiling of fPSA using the same lectins yielded much higher AUC values (0.76 and 0.80 for PHA-E and WFL, respectively). Furthermore, a combination of the GIA test based on the fPSA glycoprofile with glycoprofiling of ZAG2 did not increase the overall AUC value. Accordingly, we focused on the GIA test based on the glycoprofiling of fPSA to determine its utilisation in PCa diagnostics.

### 3.2. Potential of GIA test for therapy monitoring

In this analysis, we aimed to show the potential of the GIA test as a tool for monitoring an effect of therapy. A clear difference in glycan composition should be observed in treatment of naïve PCa patients and PCa patients under effective therapy. Fifty-nine serum samples were obtained from PCa patients who underwent therapy. In most cases (n=23), patients received hormonal therapy (sometimes in a combination with radiotherapy or chemotherapy). This set of samples was compared to a set of 109 serum samples obtained from PCa patients before they started therapy. High discrimination power for The GIA test can be confirmed by an AUC value of 0.85, which was significantly higher than the AUC value for tPSA (0.61) (**Fig. 3**). Hence, the GIA test exhibits the potential for application as a tool to monitor therapy effects. Real application of the GIA test to therapy monitoring needs to be validated in our subsequent validation study, in which a correlation will be made between results obtained from the GIA test with an accepted surrogate for therapy effectiveness, such as PSA decline or radiographic tumour regression.

**Figure 3:**
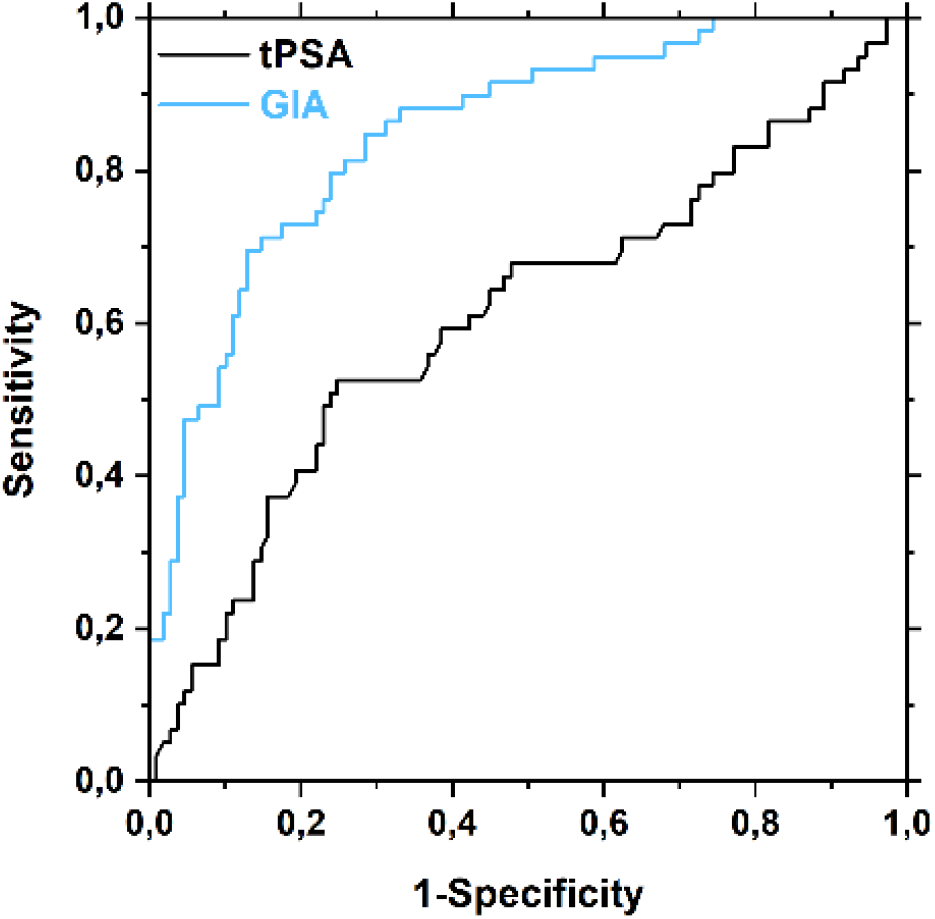
ROC analysis depicting ROC curves for tPSA (black line) and GIA test (blue line) as PCa biomarkers for therapy monitoring (PCa patients who underwent therapy *vs*. PCa patients without any treatment). Clinical validation using 168 samples revealed AUC of 0.61 for tPSA and of 0.85 for the GIA test.

### 3.3. GIA test *vs*. PHI test for PCa diagnostics

One of the second-opinion PCa serological diagnostic tests in current use is the PHI test, which combines the tPSA, fPSA and -2proPSA markers. A head-to-head study to compare the diagnostic accuracy of the PHI and GIA tests for the detection of malign and benign cases was performed using a subset of samples for which the PHI value was measured (215 serum samples in total) (**Table 1**). The AUC value was 0.69 for the PHI test and 0.81 for the GIA test, respectively (**Fig. 4**). At 95% specificity the sensitivity was 32.8% for the GIA test and 11.5% for the PHI test. At 95% sensitivity the specificity was 14.9% for the GIA test and 11.0% for the PHI test.

**Figure 4:**
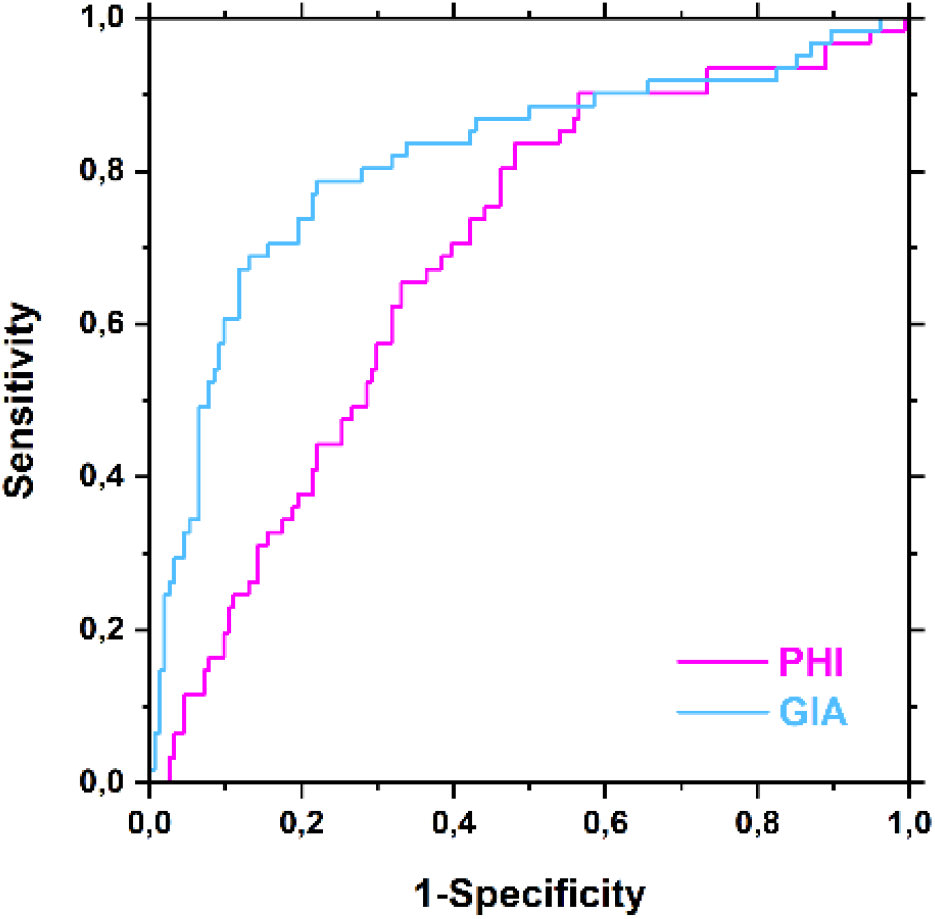
Head-to-head comparison of PHI and GIA tests: ROC analysis for PHI (magenta line) and GIA (blue line) as PCa biomarkers for early PCa diagnostics using 215 serum samples. The AUC values obtained for the PHI and GIA tests were 0.69 and 0.81, respectively.

A detailed analysis run at 80% sensitivity revealed that the following percentages of biopsies could have been identified as negative (avoidable) for the following tests: tPSA 21% of biopsies, fPSA 52% of biopsies, PHI 54% of biopsies and GIA 73% of biopsies (**Fig. 5**). The GIA test has the potential not only to significantly reduce the number of negative (avoidable) biopsies when compared with the tPSA and fPSA tests, but also when compared with an established second-opinion test, such as the PHI test. The GIA test missed five high-risk PCa cases (Gleason score ≥ 7, ISUP GG ≥2), tPSA missed four high-risk PCa cases, fPSA missed five high-risk PCa cases and fPSA% missed three high-risk PCa cases.

**Figure 5:**
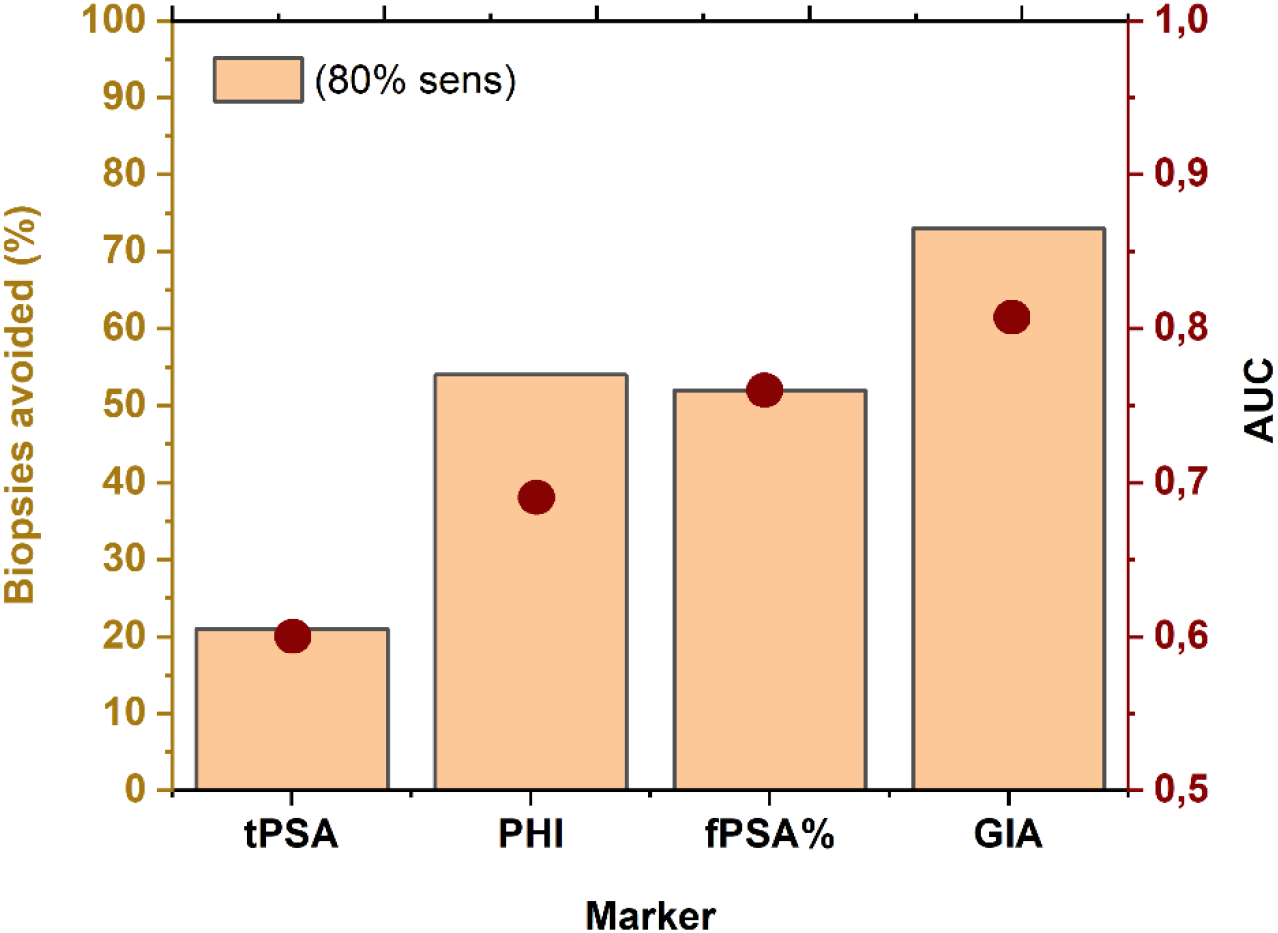
Percentage of negative (avoidable) biopsies (orange columns) calculated with 80% sensitivity for all four PCa biomarkers (tPSA, fPSA, PHI test and GIA test) from a clinical validation study performed using 215 serum samples for which PHI values were available. Negative (avoidable) biopsies were calculated as the ratio of correctly identified benign patients out of the whole BPH cohort.

### 3.4. Decision curve analysis (DCA) for GIA test

A decision curve analysis (DCA) was performed, calculating a clinical “net benefit” for diagnostic test(s) over the default strategies of diagnosing/treating all or no patients at all. Net benefit is calculated across a range of threshold probabilities (*i.e*. the minimum probability of disease at which an intervention is necessary). For the DCA, the samples of the early diagnostics cohort were used for the graph in **Fig. 6**.

**Figure 6:**
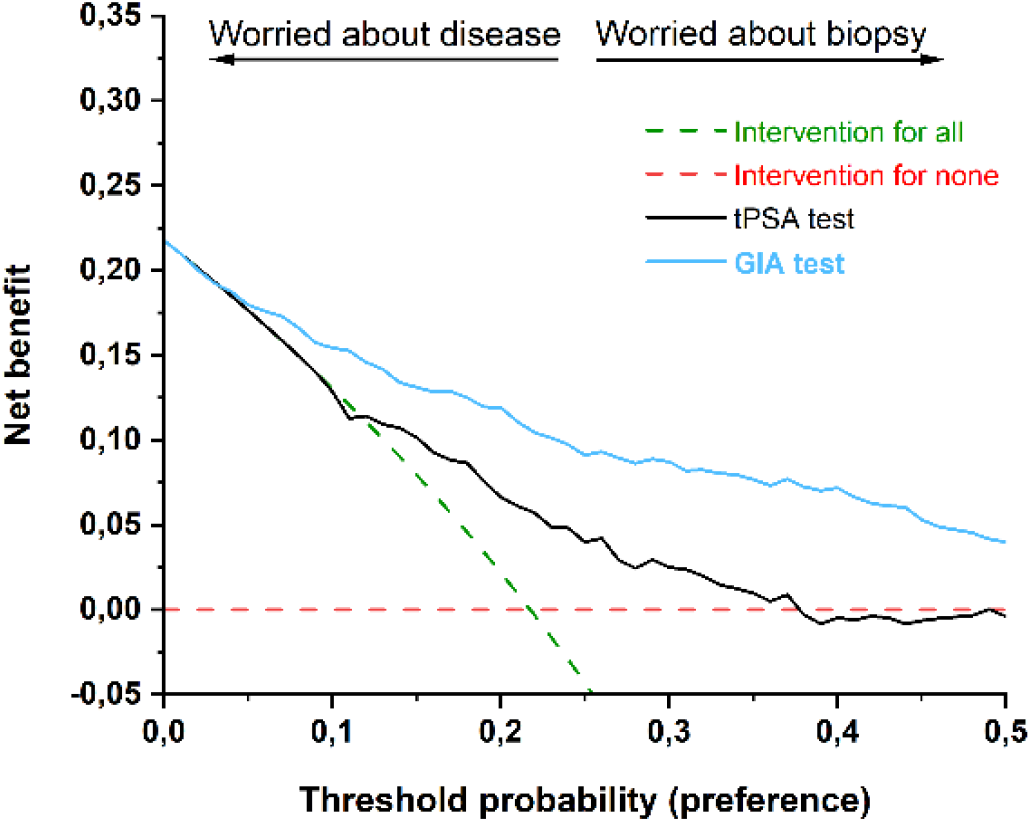
Decision curve analysis (DCA) for commonly used serological screening tPSA test (black line) and GIA test (blue line), showing two extreme strategies, *i.e*. intervention for all patients (dashed green line) and for none (dashed red line).

The main reason for using a DCA in this case is to show the potential benefit of using the GIA as a second-opinion test to determine the need for a prostate biopsy at a given threshold probability. When the probability of having a high-risk PCa is low, a urologist may decide to actively monitor the patient rather than to perform (an avoidable) biopsy. This can eliminate future barriers between patients and clinicians, as patients (not having undergone an unnecessary biopsy) will be less hesitant to return to the care-provider. On the other hand, when there is a higher probability of PCa being present, the fear is that a high-risk tumour might go undiagnosed and would subsequently be harder to cure. The DCA curve analysis indicates a net benefit of using the GIA test compared to tPSA test, since it is more efficient across all threshold probabilities, starting at ∼5% (**Fig. 6**).

### 3.5. Principal component analysis (PCA)

Principal Component Analysis (PCA) was performed on the early PCa detection cohort using OriginPro 2021b. From four different biomarkers used in the GIA test (tPSA, fPSA, fPSA^WFL^, fPSA^PHA-E^), the principal components were calculated (**Fig. 7**). The first three principal components, capturing more than 90% of the variation (line plot on the upper right), were plotted together with loading vectors in a so-called 3D biplot. Loadings (showing how strongly each parameter influences a principal component) correlating positively with PC1 are tPSA and fPSA, while fPSA^PHA-E^ and fPSA^WFL^ are correlating with PC2 and PC3, suggesting a strong added value of using these two lectins in the analysis. 95% confident ellipses are shown for benign (green) and PCa (red) cohorts (**Fig. 7**).

**Figure 7:**
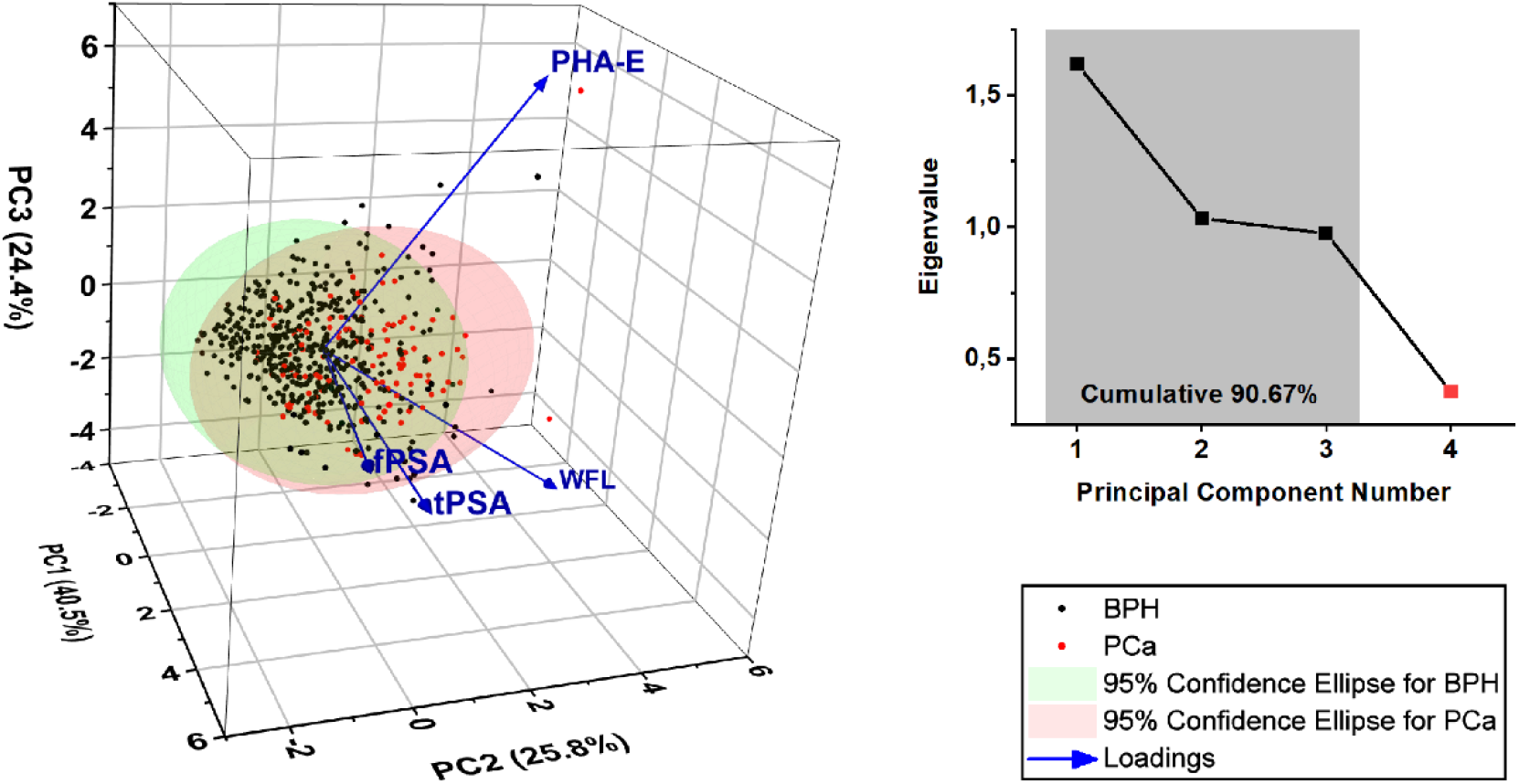
Principal component analysis (PCA) biplot showing scores and loadings for tPSA, fPSA and GIA test components (left) and eigenvalues (line plot on the right) for principal components (PC).

## 4. Discussion

PSA-based tests (serum tPSA, fPSA and PHI) are quantitative tests commonly used for PCa screening or as second-opinion tests. They are important as they may reduce overdiagnosis and overtreatment (prevent negative, avoidable biopsies). The use of different forms/precursors of PSA as PCa biomarkers is advantageous since all of them are released only by prostate tissue and thus are organ(prostate)-specific. The false negative/positive rates of the tPSA test are high, hence its use for screening or diagnostic purposes is questionable and not recommended. There is a need for identifying PCa biomarkers which are at the same time tissue- and cancer-specific.

In the present study, we confirmed that two glycoforms of fPSA recognised by WFL, (fPSA^WFL^) and PHA-E (fPSA^PHA-E^), lectins that bind to *N*-acetyl sugar residues GalNAc, LacdiNAc and bisecting GlcNAc, are highly cancer-specific, hence are useful in distinguishing malignant from benign cases, particularly in the PSA grey-zone (tPSA level in the range of 2 to 10 ng mL^-1^). The bisecting GlcNAc structure, *i.e*. a β1,4-linked GlcNAc attached to the core β-mannose residue, plays a role in tumour development and also in other physiological processes (adhesion, fertilisation, *etc*.). Since bisecting GlcNAc is not further elongated by the action of glycosyltransferases, it is considered as a special modification associated with cancer [28-30].

The same two lectins were also used for the glycoprofiling of ZA2G – an adipokine responsible for lipid mobilisation, highly expressed in cancer cachexia [31]. Unlike fPSA which usually presents with a single *N*-glycosylation site (at Asn-69), ZA2G contains 4 putative glycosylation sites (Asn-89, 92, 106 and 239), suggesting that this molecule is highly suitable for glycoprofiling in a manner similar to glycoprofiling of fPSA [32]. However, the results obtained with ZA2G glycoforms did not meet this expectation and exhibited no improvement on the discriminatory power obtained with the GIA test alone.

The DCA curve analysis, used as another statistical method independent of the ROC curve analysis, indicates a net benefit of using the GIA test compared to tPSA, as it appears to be more efficient across all threshold probabilities, starting at ∼5%. PCA also suggests a strong added value of using lectins for PCa diagnostics. It is worth mentioning that both models showed a potential gain in comparison with the “biopsy-all” and “biopsy-none” models across the strategy thresholds. A substantial gain was achieved for the GIA test over routine PSA screening. An observation made by DCA analysis was also confirmed by ROC analysis, when the addition of glycoprofiling of fPSA by two lectins in the GIA test significantly improved the AUC value of the combination of tPSA and fPSA alone from 0.78 to 0.84.

The clinical utility of the GIA test was compared to other tests applied to PCa screening/diagnostics or as second-opinion tests to determine which men should be biopsied. At 80% sensitivity, the GIA test identified 70-73% of the biopsies as negative (and thus avoidable), whereas other tests identified a lower proportion of biopsies as negative (avoidable), *i.e*. tPSA (21-30%), fPSA (52-53%) and the PHI test (54%) (**Figs. 2** and **5**). The results here showed that the GIA test outperformed other PSA-based quantitative tests in terms of its AUC value and avoidable biopsies count. Choosing the right lectins for the glycoprofiling of proteins is crucial for this kind of assay. Only certain glycan patterns would appear to carry the cancer-specific biological information caused by fragmentation of the Golgi apparatus in cancer cells [33].

The GIA test has the potential to be used in therapy-monitoring since it distinguished between treated and untreated PCa patients significantly better (AUC value of 0.85) than tPSA (AUC value of 0.61 (**Fig. 3**)). The rationale behind using the GIA test for therapy-monitoring is that therapy induces a decrease in the number of cancerous cells. Hence, as a result, the level of glycans associated with cancer is also decreasing and the glycan-modifying enzymes synthesise “healthy” glycans [34, 35]. Thus, the glycans produced by patients responding to a therapy resemble the glycans of healthy individuals. Our results warrant further evaluation of this application of the GIA test. Any test which can indicate effective treatment would be very helpful in suppressing negative effects associated with the disease, since PCa patients can suffer from PSAdynia and experience psychological anxiety or problems in relationships [36, 37].

The gold standard in glycan analysis is still instrumental-based, integrating mass spectrometry with separation methods [38]. Such an approach is, however, costly and time-consuming, requiring a lengthy data-processing and assessment procedure, hence it is hardly compatible with clinical practice. Lectin-based approaches, on the other hand, can be effectively applied to glycan analysis in ELISA-like formats, which are fully compatible with clinical practice [39]. A further advantage of using lectins is the possibility of deploying them for glycan analysis in complex samples without any pre-treatment and for the glycoprofiling of intact proteins [39]. The GIA test based on the integration of modified magnetic beads, as used in this study, overcomes the challenges typical of lectin-assisted glycoprofiling of proteins [20], while affording the possibility of working in an ELISA-like format that is available in any routine clinical laboratory.

## 5. Conclusions

The clinical validations revealed that the glycoprofiling of ZA2G showed little potential for PCa diagnostics, while the glycoprofiling of fPSA was of significant clinical potential. The GIA test integrating glycoprofiling of fPSA (fPSA^WFL^ and fPSA^PHA-E^) could be used in PCa early diagnostics (AUC=0.84; n=501 samples) and in discriminating between therapy-naïve PCa patients and patients in therapy (AUC=0.85; n=168 samples). Moreover, the GIA test (AUC=0.81) outperformed the PHI test (AUC=0.69) in early diagnostics in a head-to-head comparison run on a subset of serum samples (n=215 samples). Furthermore, out of 392 negative biopsies considered to be avoidable, 70-73% could have been prevented had the GIA test been used; 21-30% had the tPSA been used; 52-53% with use of the fPSA and 54% with use of the PHI test. Accordingly, the GIA test is able to outperform all the PSA-based serological tests and has the ability to significantly reduce the number of biopsies.

## 6. Acknowledgement

The financial support received from the Slovak Research and Development Agency APVV-21-0329 and APVV*-*20-0476 is gratefully acknowledged. The publication was supported by EIC Accelerator grant 190185443 (HORIZON). This study was supported by the Ministry of Health of the Czech Republic - conceptual development of research organisation (Faculty Hospital in Pilsen - FNPl, 00669806), BBMRI-CZ: Biobank network - a versatile platform for research of the etiopathogenesis of diseases CZ.02.1.01/0.0/0.0/16_013/000167 and LM2015089, and by the Cooperation Programme, research area Pharmaceutical Sciences. The authors would like to express their sincere gratitude to Jana Trpkova, Andrea Eigentler and Gabriele Dobler for handling the serum samples used in the study, and to Eberhard Steiner for preparation of the clinical data.

## Notes

### Competing Interest Statement

The authors have declared no competing interest.

## References

1. Sung H, Ferlay J, Siegel RL, Laversanne M, Soerjomataram I, Jemal A, et al. Global cancer statistics 2020: GLOBOCAN estimates of incidence and mortality worldwide for 36 cancers in 185 countries. CA: Cancer J Clin. 2021;71(3):209–49.

2. Wright P, Wilding S, Watson E, Downing A, Selby P, Hounsome L, et al. Key factors associated with social distress after prostate cancer: Results from the United Kingdom Life after Prostate Cancer diagnosis study. Cancer Epidemiol. 2019;60:201–7.

3. Houédé N, Rébillard X, Bouvet S, Kabani S, Fabbro-Peray P, Trétarre B, et al. Impact on quality of life 3 years after diagnosis of prostate cancer patients below 75 at diagnosis: an observational case-control study. BMC Cancer. 2020;20(1):1–12.

4. Trujillo B, Wu A, Wetterskog D, Attard G. Blood-based liquid biopsies for prostate cancer: clinical opportunities and challenges. Br J Cancer. 2022;127(8):1394–402.

5. Heijnsdijk EA, de Carvalho TM, Auvinen A, Zappa M, Nelen V, Kwiatkowski M, et al. Cost-effectiveness of prostate cancer screening: a simulation study based on ERSPC data. J Natl Cancer Inst. 2015;107(1):366.

6. Vickers AJ. Redesigning prostate cancer screening strategies to reduce overdiagnosis. Clin Chem. 2019;65(1):39–41.

7. Campos-Fernández E, Barcelos LS, de Souza AG, Goulart LR, Alonso-Goulart V. Research landscape of liquid biopsies in prostate cancer. Am J Cancer Res. 2019;9(7):1309.

8. Bai Y, Zhao H. Liquid biopsy in tumors: opportunities and challenges. Annals Translat Med. 2018;6(Suppl 1):S89.

9. Bertok T, Bertokova A, Hroncekova S, Chocholova E, Svecova N, Lorencova L, et al. Novel prostate cancer biomarkers: Aetiology, clinical performance and sensing applications. Chemosensors. 2021;9(8):205.

10. Bertokova A, Svecova N, Kozics K, Gabelova A, Vikartovska A, Jane E, et al. Exosomes from prostate cancer cell lines: Isolation optimisation and characterisation. Biomed Pharmacother. 2022;151:113093.

11. Tkac J, Bertok T, Hires M, Jane E, Lorencova L, Kasak P. Glycomics of prostate cancer: Updates. Exp Rev Proteomics. 2019;16(1):65–76.

12. Tkac J, Gajdosova V, Hroncekova S, Bertok T, Hires M, Jane E, et al. Prostate-specific antigen glycoprofiling as diagnostic and prognostic biomarker of prostate cancer. Interface Focus. 2019;9(2):20180077.

13. Petrosyan A. Onco-Golgi: is fragmentation a gate to cancer progression? Biochem Mol Biol J. 2015;1(1):16.

14. Bui S, Mejia I, Díaz B, Wang Y. Adaptation of the Golgi apparatus in cancer cell invasion and metastasis. Front Cell Develop Biol. 2021;9:806482.

15. Zhang X. Alterations of golgi structural proteins and glycosylation defects in cancer. Front Cell Develop Biol. 2021;9:665289.

16. Liu L, Doray B, Kornfeld S. Recycling of Golgi glycosyltransferases requires direct binding to coatomer. Proc Natl Acad Sci USA. 2018;115(36):8984–9.

17. Tu L, Banfield DK. Localization of Golgi-resident glycosyltransferases. Cell Mol Life Sci. 2010;67:29–41.

18. Bertok T, Jane E, Bertokova A, Lorencova L, Zvara P, Smolkova B, et al. Validating fPSA glycoprofile as a prostate cancer biomarker to avoid unnecessary biopsies and re-biopsies. Cancers. 2020;12(10):2988.

19. Bertokova A, Bertok T, Jane E, Hires M, Ďubjaková P, Novotná O, et al. Detection of N, N-diacetyllactosamine (LacdiNAc) containing free prostate-specific antigen for early stage prostate cancer diagnostics and for identification of castration-resistant prostate cancer patients. Biorg Med Chem. 2021;39:116156.

20. Bertok T, Tkac J, inventors; PCT/EP2019/057386. https://patentscope.wipo.int/search/en/detail.jsf?docId=WO2019185515, assignee. Means and methods for glycoprofiling of a protein 2021.

21. Bertok T, Jane E, Bertokova A, Lorencova L, Zvara P, Smolkova B, et al. Validating fPSA Glycoprofile as a Prostate Cancer Biomarker to Avoid Unnecessary Biopsies and Re-Biopsies. Cancers. 2020;12(10). doi: 10.3390/cancers12102988. PubMed PMID: WOS:000584073600001.

22. Bertokova A, Bertok T, Jane E, Hires M, Ďubjaková P, Novotná O, et al. Detection of N, N-diacetyllactosamine (LacdiNAc) containing free prostate-specific antigen for early stage prostate cancer diagnostics and for identification of castration-resistant prostate cancer patients. Biorg Med Chem. 2021:116156.

23. Peracaula R, Tabarés G, Royle L, Harvey DJ, Dwek RA, Rudd PM, et al. Altered glycosylation pattern allows the distinction between prostate-specific antigen (PSA) from normal and tumor origins. Glycobiology. 2003;13(6):457–70. doi: 10.1093/glycob/cwg041.

24. Pihikova D, Pakanova Z, Nemcovic M, Barath P, Belicky S, Bertok T, et al. Sweet characterisation of prostate specific antigen using electrochemical lectin-based immunosensor assay and MALDI TOF/TOF analysis: Focus on sialic acid. Proteomics. 2016;16(24):3085–95. doi: 10.1002/pmic.201500463. PubMed PMID: WOS:000390809000006.

25. Pihíková D, Belicky Š, Kasák P, Bertok T, Tkac J. Sensitive detection and glycoprofiling of a prostate specific antigen using impedimetric assays. Analyst. 2016;141(3):1044–51. doi: 10.1039/C5AN02322J.

26. Tkac J, Bertok T, inventors; PCT/EP2022/072138, https://patentscope.wipo.int/search/en/detail.jsf?docId=WO2023012352&_cid=P22-LDXXGC-67374-1, assignee. Standard for glycoprofiling of proteins 2023.

27. Bertok T, Bertokova A, Jane E, Hires M, Aguedo J, Potocarova M, et al. Identification of whole-serum glycobiomarkers for colorectal carcinoma using reverse-phase lectin microarray. Front Oncol. 2021;11:735338.

28. Nyalwidhe JO, Betesh LR, Powers TW, Jones EE, White KY, Burch TC, et al. Increased bisecting N-acetylglucosamine and decreased branched chain glycans of N-linked glycoproteins in expressed prostatic secretions associated with prostate cancer progression. Proteom Clin Appl. 2013;7(9-10):677–89.

29. Kohler RS, Anugraham M, López MN, Xiao C, Schoetzau A, Hettich T, et al. Epigenetic activation of MGAT3 and corresponding bisecting GlcNAc shortens the survival of cancer patients. Oncotarget. 2016;7(32):51674–86.

30. Chen Q, Tan Z, Guan F, Ren Y. The essential functions and detection of bisecting GlcNAc in cell biology. Front Chem. 2020;8:511.

31. Hassan MI, Waheed A, Yadav S, Singh TP, Ahmad F. Zinc α2-glycoprotein: a multidisciplinary protein. Mol Cancer Res. 2008;6(6):892–906.

32. Butler W, Huang J. Glycosylation Changes in Prostate Cancer Progression. Front Oncol. 2021;11:809170. doi: 10.3389/fonc.2021.809170.

33. Bajaj R, Warner AN, Fradette JF, Gibbons DL. Dance of The Golgi: Understanding Golgi Dynamics in Cancer Metastasis. Cells. 2022;11(9):1484. Epub 2022/05/15. doi: 10.3390/cells11091484. PubMed PMID: 35563790; PubMed Central PMCID: PMCPMC9102947.

34. Narimatsu Y, Joshi HJ, Nason R, Van Coillie J, Karlsson R, Sun L, et al. An Atlas of Human Glycosylation Pathways Enables Display of the Human Glycome by Gene Engineered Cells. Molecular Cell. 2019;75(2):394-407.e5. doi: https://doi.org/10.1016/j.molcel.2019.05.017.

35. Narimatsu Y, Büll C, Chen Y-H, Wandall HH, Yang Z, Clausen H. Genetic glycoengineering in mammalian cells. J Biol Chem. 2021;296:100448. doi: https://doi.org/10.1016/j.jbc.2021.100448.

36. Mathew S, Rapsey CM, Wibowo E. Psychosocial Barriers and Enablers for Prostate Cancer Patients in Starting a Relationship. J Sex Marital Ther. 2020;46(8):736–46. Epub 2020/08/25. doi: 10.1080/0092623x.2020.1808549. PubMed PMID: 32835628.

37. Klotz LH. PSAdynia and other PSA-related syndromes: a new epidemic--a case history and taxonomy. Urology. 1997;50(6):831–2. Epub 1998/01/14. doi: 10.1016/s0090-4295(97)00490-1. PubMed PMID: 9426708.

38. Pihikova D, Kasak P, Kubanikova P, Sokol R, Tkac J. Aberrant sialylation of a prostate-specific antigen: Electrochemical label-free glycoprofiling in prostate cancer serum samples. Anal Chim Acta. 2016;934(-):72–9. Epub 2016/08/11. doi: 10.1016/j.aca.2016.06.043. PubMed PMID: 27506346; PubMed Central PMCID: PMCPMC5659379.

39. Paleček E, Tkáč J, Bartosik M, Bertók Ts, Ostatná V, Paleček J. Electrochemistry of nonconjugated proteins and glycoproteins. Toward sensors for biomedicine and glycomics. Chem Rev. 2015;115(5):2045–108.

